# Myeloperoxidase-induced fibrinogen unfolding and clotting

**DOI:** 10.1101/2021.01.06.425586

**Authors:** Nikolay A. Barinov, Elizaveta R. Pavlova, Anna P. Tolstova, Evgeniy V. Dubrovin, Dmitry V. Klinov

## Abstract

Fibrinogen is a major protein of blood coagulation system and is a promising component of biomaterials and protein matrixes. Conformational changes of fibrinogen underlie the important mechanism of thrombin mediated fibrinogen clotting but also may induce the loss of its biological activity and (amyloid) aggregation. Understanding and controlling of fibrinogen unfolding is important for the development of fibrinogen based materials with tunable properties. We have discovered that myeloperoxidase induces denaturation of fibrinogen molecules followed by fibrinogen clotting, which is not thrombin-dependent. This is the first example of ATP-independent, non-targeted protein-induced protein denaturation. The morphological structure of unfolded fibrinogen molecules and “non-conventional” fibrinogen clots has been characterized using high-resolution atomic force microscopy and scanning electron microscopy techniques. Circular dichroism (CD) spectroscopy has shown no significant changes of the secondary structure of the fibrinogen clots. The absorbance spectrophotometry has demonstrated that the kinetics of myeloperoxidase induced fibrinogen clotting strongly decays with growth of ionic strength indicating a major role of the Debye screening effect in regulating of this process. The obtained results provide with the novel concepts of protein unfolding and open new insights into fibrinogen clotting. Moreover, they give new possibilities in biotechnological and biomedical applications, e.g., for regulation of fibrinogen clotting and platelet adhesion and for the development of fibrinogen-based matrices.

The organization of a protein molecule is characterized by different hierarchical levels such as primary, secondary, tertiary and quaternary structure. Protein unfolding or denaturation, i.e. its transformation to a lower order structure (and loss of a higher order structure), is a biologically and biotechnologically relevant process. Protein unfolding is a prerequisite for an alternative folding pathway including amyloid aggregation ^1,2^. Unfolded proteins may be used in development of protein films and coatings with special properties such as enhanced mechanical stability ^3–5^, resistance to protein adsorption or platelet adhesion ^6,7^ and other advantages ^8^. Unfolding of a protein molecule may lead to the loss of its biological function ^9^ that has important consequences in biosensor ^10,11^ and pharmaceutical applications ^12^.

---

The known chemical substances and physicochemical factors that may induce protein unfolding include urea, organic acids, alcohols, different salts, detergents, change of pH, heating, pressure, interaction with surfaces, etc. ^12^ To the best of our knowledge, ATP-independent, non-targeted interaction with other proteins has not been reported to induce protein unfolding. However, protein may be considered as an appealing denaturing agent in biomedical applications due to its high biocompatibility and nontoxicity. The aim of this work was to reveal the effect of protein unfolding caused by its contact with another protein. Two medically relevant proteins of blood plasma such as an anionic 340 kDa fibrinogen and a 140 kDa heme-containing homodimeric globular cationic enzyme myeloperoxidase have been used as model proteins.

Fibrinogen is one of the most abundant proteins of the blood plasma and is a key protein of blood coagulation system. Fibrinogen is composed of three pairs of polypeptide chains held together by disulfide bonds ^13^. Fibrinogen polypeptide chains are organized in three globular regions (one central and two outer ones) connected by coiled coils and contain two unstructured αC regions. Though fibrinogen is an anionic protein (pI ∼5.8 ^14^), it has a heterogeneous charge distribution: fibrinogen globular regions are negatively charged whereas αC regions are positively charged at neutral pH ^15^.

Myeloperoxidase is secreted by activated neutrophilic leukocytes into phagosomes and extracellular space. To protect the body from invading pathogens myeloperoxidase catalyzes the production of hypochlorous acid and other oxidizing agents^16^. Myeloperoxidase (pI ∼10.7) has been shown to bind with anionic proteins and negatively charged cell surfaces via electrostatic interaction.^17^

Fibrinogen was shown to bind with the proteins present in blood plasma leading to important physiological reactions such as serum amyloid A induced promotion of fibrin amyloid formation ^18^ or β-amyloid induced blockage of plasmin-mediated cleavage ^19^. Fibrinogen conformation was shown to mediate blood clotting, platelet adhesion ^20–22^, inflammation ^23,24^, and endothelial cell proliferation ^25^, therefore, controlled fibrinogen unfolding can potentially regulate these important processes. In particular, myeloperoxidase-induced fibrinogen unfolding and its consequences on fibrinogen clotting may be of great biotechnological and medical interest.

Recently we have visualized unfolding of individual fibrinogen molecules induced by their contact with a highly oriented graphite (HOPG) surface modified with N,N’-(decane-1,10-diyl)bis(tetraglycineamide) (GM) ([Gly_4_-NHCH_2_]_2_C_10_H_20_)^26,27^. We have also developed an approach of AFM characterization of unfolding of individual protein molecules induced by the factors other than contact with a substrate surface. This approach implies protein deposition on a GM-HOPG surface followed by immediate drying.

The developed approach has been used in this work to characterize fibrinogen interaction with myeloperoxidase at a single-molecule level. Moreover, influence of myeloperoxidase on fibrinogen clotting has been investigated using SEM, absorbance spectrophotometry and circular dichroism spectroscopy. The obtained results demonstrate the first example of non-targeted protein-induced protein unfolding and a novel pathway of fibrinogen clotting.

## Results and Discussions

### AFM characterization of fibrinogen unfolding by myeloperoxidase

Fibrinogen molecules adsorbed onto a GM-HOPG surface for 5 s are characterized by a trinodular structure with two outer globular regions 3 – 3.5 nm high and one central globular region ∼2 nm high ^27–29^. Myeloperoxidase molecules deposited by the same procedure adsorb onto a GM-HOPG surface as globular structures ∼4 nm high ^30^. The AFM images of 1:1 (mass ratio) mixture of fibrinogen with myeloperoxidase incubated for 3 s at 21°C and deposited onto a GM-HOPG surface (Figure 1a; three regions marked with the boxes are enlarged on the right) have revealed the presence of typical for myeloperoxidase molecules single globules (some examples are marked with arrows) and typical for fibrinogen molecules trinodular structures (e.g., regions I–III), however, with different size of outer globules. Three types of trinodular structures can be distinguished depending on a height of the outer globules (see the height profiles in Figure 1b along the dotted lines in the enlarged regions of AFM images): the structures with the outer globules 3 – 3.5 nm in height (e.g., region III) can be assigned to fibrinogen molecules, which haven’t formed complexes with myeloperoxidase, whereas the structures with significantly higher one (e.g., region I) or both (e.g., region II) outer globules – to the fibrinogen–myeloperoxidase complexes with 1:1 or 1:2 fibrinogen–myeloperoxidase molar ratio, respectively. The height distribution of the globules of trinodular structures (Figure 1c) demonstrates the peaks around 2 and 3.2 nm that correspond to inner and outer globules of native-like fibrinogen molecules (e.g., region III) respectively ^27–29^. Moreover, it has one more maximum at ∼4 nm that we associate with fibrinogen–myeloperoxidase complexes. The presence of larger height values in the distribution (up to 6 nm) may be connected with the adsorption of the asymmetrical globular structures of fibrinogen–myeloperoxidase complexes on a substrate by their different sides. Importantly that fibrinogen–myeloperoxidase complexes form quite fast (within 3 s after mixing the proteins), and each outer globular region of a fibrinogen molecule may independently bind a myeloperoxidase molecule.

**Figure 1.**
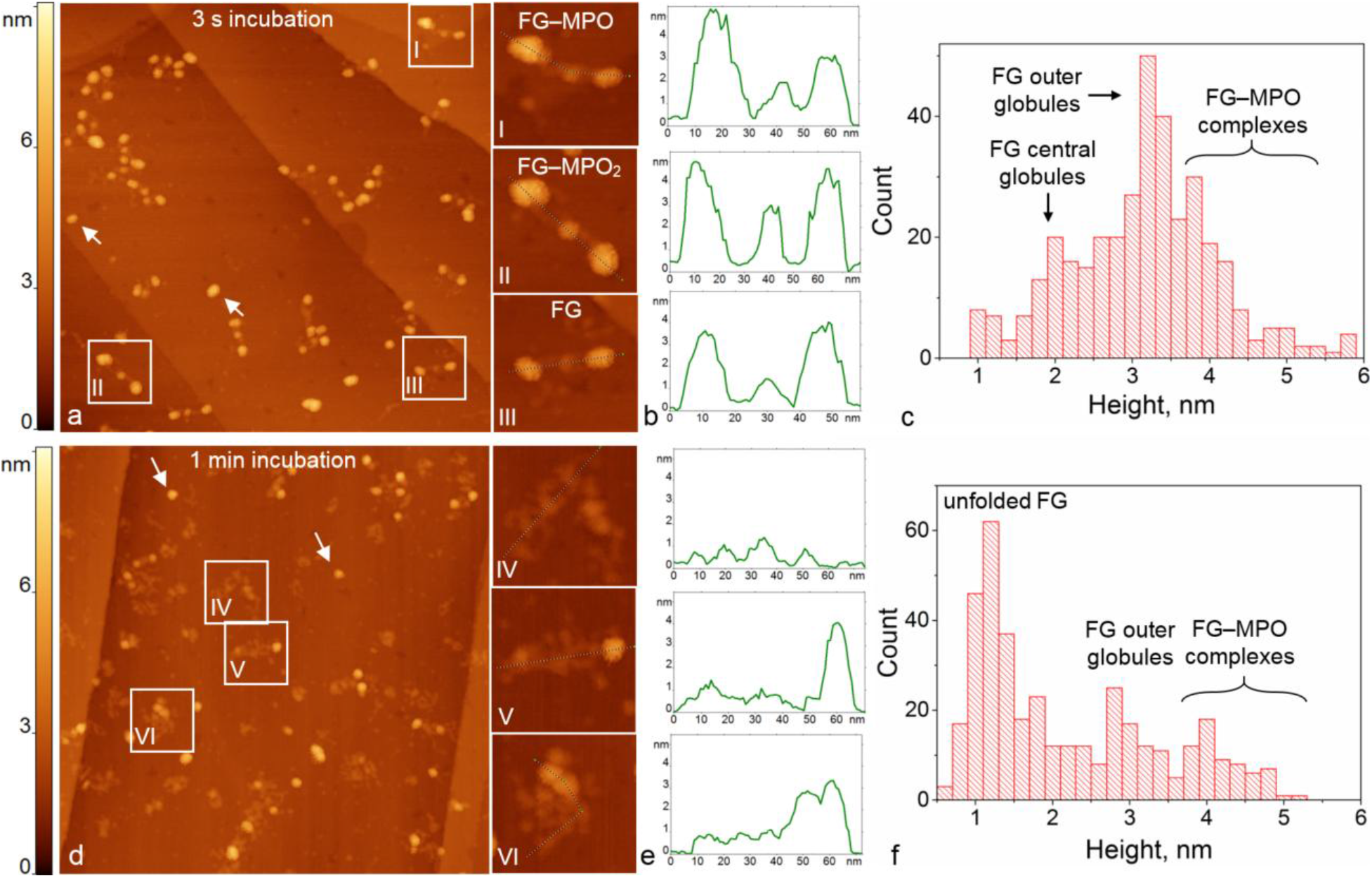
Visualization of fibrinogen (FG)–myeloperoxidase (MPO) complexes at a single-molecule level. (a,d) AFM height images of fibrinogen fibrinogen–myeloperoxidase mixture incubated for (a) 3 s and (d) 1 min, and deposited onto a GM-HOPG surface. The regions marked with a box and labelled with Roman numbers are enlarged on the right. Individual myeloperoxidase molecules are marked with arrows. The sizes of the images are 500 × 500 nm^2^ (enlarged regions 70 × 70 nm^2^). (b), (e) Height profiles along the dotted lines in the corresponding regions of the AFM images. (c), (f) Height distribution histograms of the objects observed in the corresponding AFM images (N=364 for (c) and N=382 for (f)).

The electrostatics was shown to play a primary role in fibrinogen interaction with the charged surfaces. The extended and anisotropic charge distribution of a fibrinogen molecule allows fibrinogen adsorption even on likely charged surfaces ^14,31^. Therefore, we assume that the observed fast formation of fibrinogen–myeloperoxidase complex in water also has the electrostatic nature. Under the conditions of the experiment (pH of deionized water 5.5) myeloperoxidase (pI>10 ^17^) is highly positively charged, whereas fibrinogen is approximately neutral. However, fibrinogen molecules contain negatively charged patches in the outer globular regions (Supplementary information, Figure S1), which can interact with myeloperoxidase molecules in the absence of screening charges.

In contrast, fibrinogen does not form molecular complexes with another globular blood plasma protein, ferritin, even after 30 minutes of their incubation in water: the AFM images of the fibrinogen–ferritin mixture demonstrate these proteins separately (Supplementary information, Figure S2). This observation can be explained by the fact that ferritin (pI ∼5.2–5.7 ^32^) is almost neutral at pH 5.5 and therefore does not interact electrostatically with fibrinogen.

An increase of the incubation time of fibrinogen with myeloperoxidase to 1 min has led to dramatic differences in the morphology of fibrinogen–myeloperoxidase complexes: many fibrillary structures appear in AFM images (Figure 1d; three regions marked with the boxes are enlarged on the right). The height profiles along the dotted lines in enlarged regions of AFM images are shown in Figure 1e. The height of the fibrils is about 1.2 nm (the most pronounced maximum in the height distribution histogram in Figure 1f) indicating significant decrease as compared to fibrinogen molecules and fibrinogen–myeloperoxidase complexes (Figure 1c). The fibrinogen–myeloperoxidase complexes with trinodular structure are almost absent in the AFM images. The observed transformation of fibrinogen and fibrinogen–myeloperoxidase complexes from trinodular (Figure 1a) to predominantly fibrillary (Figure 1d) structures can be associated with fibrinogen unfolding (denaturation), which is induced by a fibrinogen contact with myeloperoxidase molecules. Similar unfolding of fibrinogen conformation has been observed after several minutes of fibrinogen adsorption on a GM-HOPG surface ^26,27^. However, in the present work the contribution of surface-induced unfolding of fibrinogen molecules is negligible due to relatively small time of adsorption (5 s) ^26–29^.

Apart from fibrillary structures AFM images of fibrinogen–myeloperoxidase mixture incubated for 1 min contain also globular features 3–4 nm in height (Figure 1d and two smaller maximums in height distribution in Figure 1f). While a ∼3 nm maximum may be associated with outer globular regions of fibrinogen molecules that did not form complexes with myeloperoxidase and, therefore, did not unfold (e.g. in region VI), a ∼4 nm maximum may correspond to myeloperoxidase molecules (e.g., region V). Some of them are located within the unfolded fibrinogen molecules (e.g., in region V), however, isolated myeloperoxidase molecules are also present in the AFM images (see the arrows in Figure 1d). This fact together with the observation of purely fibrillary structures testifies that fibrinogen unfolding induces dissociation of myeloperoxidase molecules from the complexes. This dissociation may take place due to the change of spatial charge distribution of a fibrinogen molecule caused by its reorientation upon unfolding.

### Characterization of myeloperoxidase-induced fibrinogen fibers

It is well-known that thrombin induces proteolytic cleavage of fibrinogen molecules and formation of fibrin fibers ^13^. Typical AFM image of an oligomer fibrin fiber formed in 10 min after thrombin addition to a fibrinogen solution is presented in Figure 2a; it consists of the periodically following oval and “heart-shaped” structures 3–4 nm in height (e.g., Figure 2a, bottom), in accordance with the previously reported results ^28,33^. Similar fibers are observed in 10 min after addition of thrombin to a 1 min mixture of fibrinogen with myeloperoxidase (Figure 2b, arrows). However, these fibers are accompanied by an extensive network of thinner (∼1 nm in height) fibers (Figure 2b, bottom). Analogous thin fibrillary networks have been formed in a fibrinogen–myeloperoxidase mixture incubated for 10 min (Figure 2c) without thrombin. Therefore, we assume that upon addition of thrombin into myeloperoxidase–fibrinogen mixture fibrinogen has two pathways for aggregation: conventional fibrinogen clotting takes place with a native-like fraction of fibrinogen molecules upon addition of thrombin, whereas a myeloperoxidase unfolded fraction of fibrinogen molecules aggregate in an extensive fibrillary network composed of thin fibrils. The latter process is not mediated by thrombin.

**Figure 2.**
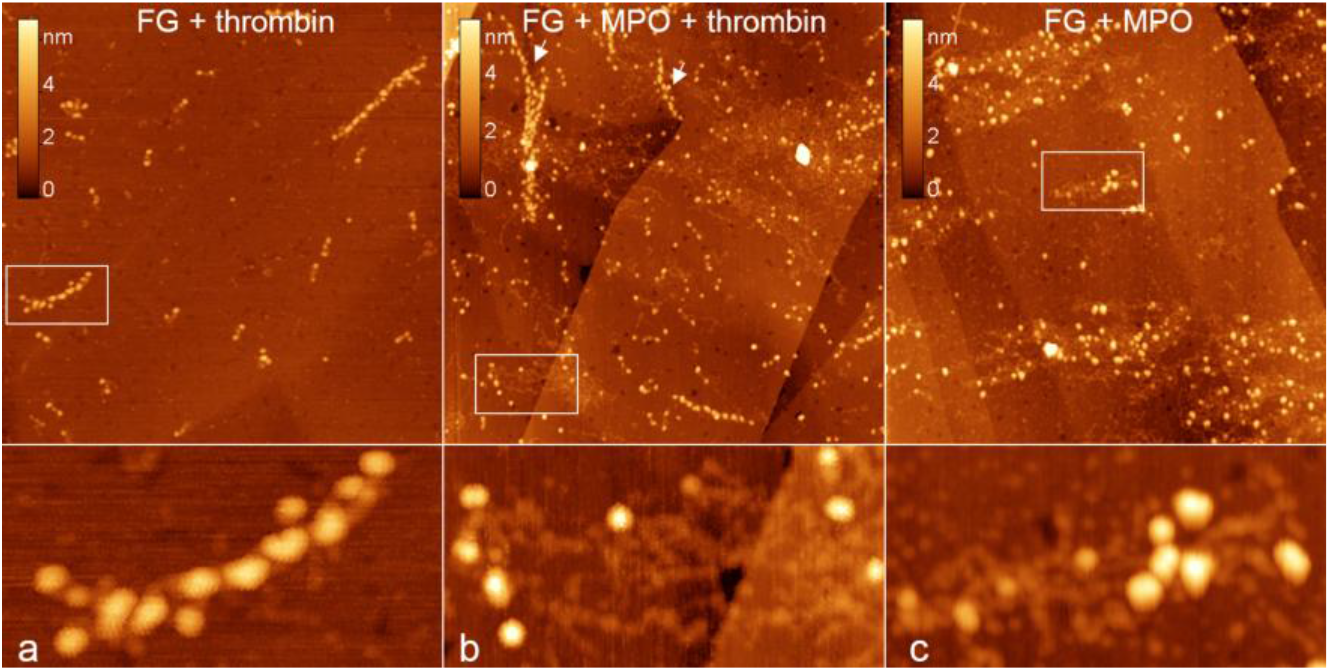
AFM images of fibrinogen fibers. AFM height images of the fibrils formed after (a) fibrinogen incubation with thrombin for 10 min, (b) fibrinogen incubation with myeloperoxidase for 1 min followed by addition of thrombin for 10 min (fibrin fibers are marked with arrows), (c) fibrinogen incubation with myeloperoxidase for 10 min. The regions marked with a box are enlarged at the bottom. The sizes of the images are 1000 × 1000 nm^2^ (enlarged regions 200 × 100 nm^2^).

Furthermore, using SEM we have characterized at a microscale the structures formed by myeloperoxidase unfolded fibrinogen at a higher fibrinogen concentration (150 µg/ml; Figure 3a): they consist of the bundles of relatively short (1–5 µm) fibers 100–200 nm in diameter. These fibers aggregate quite densely so that most interstices are less than 500 nm; moreover, they form even denser areas, which are almost devoid of interstices (arrow) and have the form of swelling.

**Figure 3.**
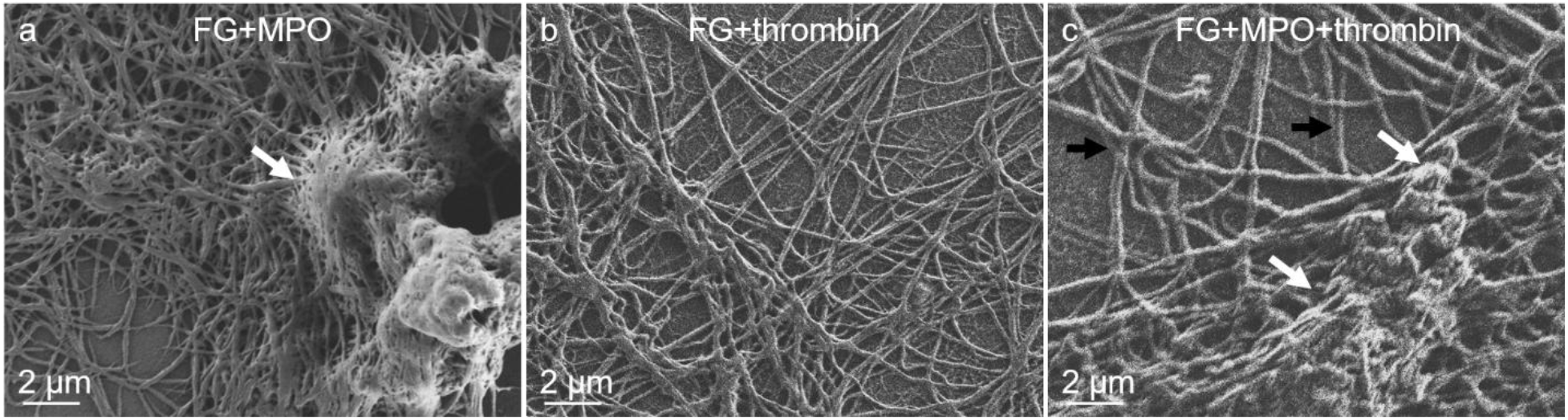
SEM images of fibrinogen fibrils. SEM images of fibrinogen fibrils formed after (a) fibrinogen incubation with myeloperoxidase for 1 min, (b) fibrinogen incubation with thrombin for 1 min, (c) fibrinogen incubation with myeloperoxidase for 1 min followed by addition of thrombin for 1 min.

A typical SEM image of the conventional (thrombin mediated) fibrin clot in the solution with the same fibrinogen concentration is presented in Figure 3b. It may be characterized by considerably longer fibrils (tens of microns) and larger (several microns) interstices that is in general agreement with the previously published results ^34^.

The structures formed after thrombin addition to a myeloperoxidase–fibrinogen mixture contain both types of fibrils described above (Figure 3c): relatively short and densely packed fibrils (white arrows) are accompanied by longer and less densely organized ones (black arrows). This observation allows us to conclude that both types of fibrinogen clotting take place: non-conventional clotting of myeloperoxidase unfolded fibrinogen and conventional (thrombin mediated) clotting of the fraction of native fibrinogen molecules, which have not interacted with myeloperoxidase molecules.

### Spectrophotometric characterization of fibrinogen interaction with myeloperoxidase in water

For in situ characterization of fibrinogen fibril formation we have used absorption spectrophotometry. Two bottom absorption spectra in Figure 4a refer to the fibrinogen solution before (bottom curve) and 30 minutes after (second bottom curve) addition of myeloperoxidase. Non-zero signal in the latter spectrum at the wavelengths larger than 320 nm originates from light scattering that is indicative of protein aggregation ^35^. Increase of the light absorption signal of the fibrinogen–myeloperoxidase mixture in the “near-UV” region (240–300 nm) can be attributed both to light scattering by the fibrillary protein aggregates and to the increase of the number of chromophores due to addition of myeloperoxidase solution. However, the latter factor is quite weak due to a 50-fold lower concentration of myeloperoxidase than fibrinogen. After addition of thrombin to a 30 min fibrinogen–myeloperoxidase mixture the absorption spectrum of the sample becomes significantly higher (third bottom curve in Figure 4a) indicating further increase of protein aggregation.

**Figure 4.**
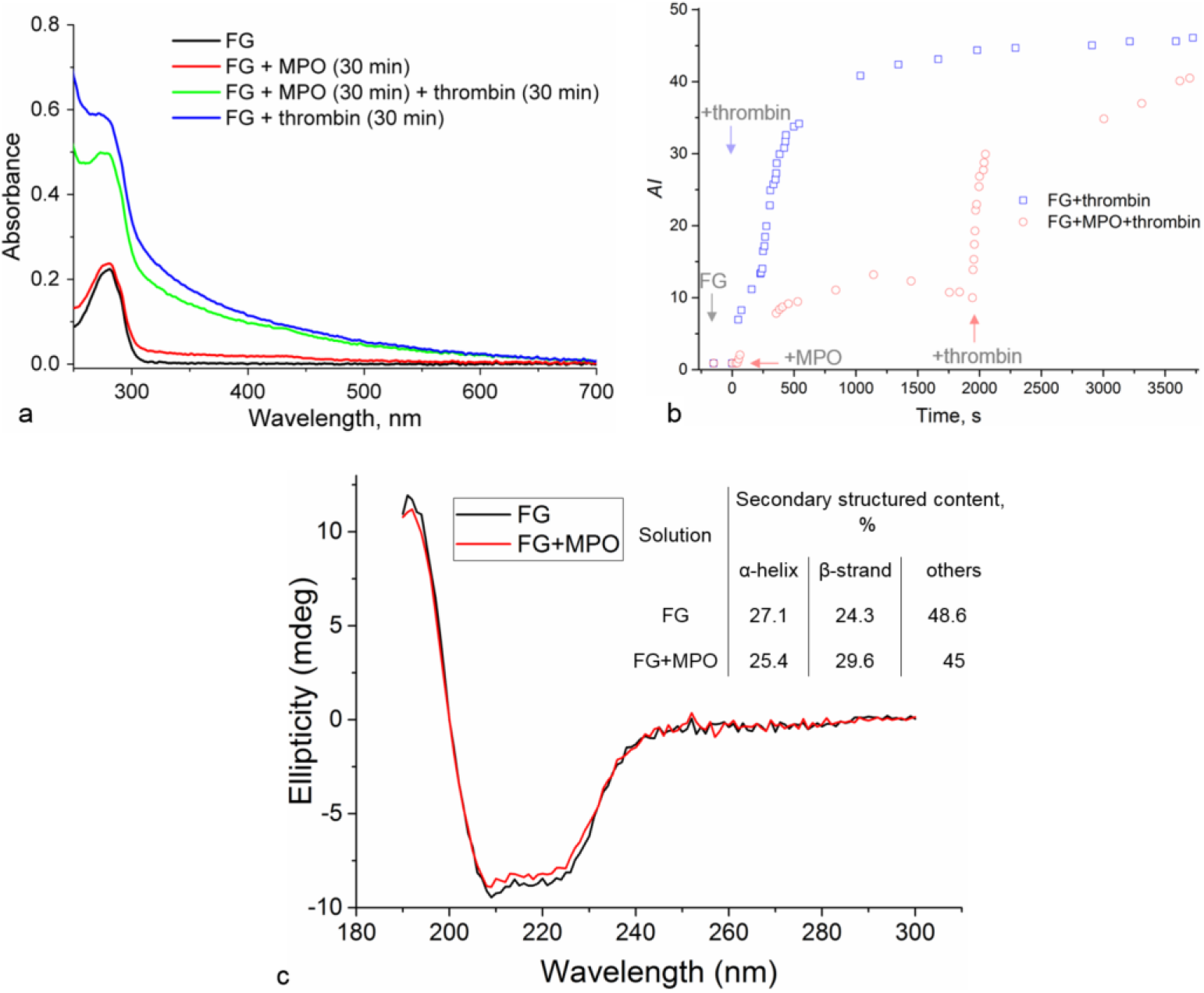
Absorbance spectrophotometry and CD spectroscopy of fibrinogen fibrils. (a) Light absorbance spectra of the solutions of fibrinogen and fibrinogen incubated with other proteins. (b) Time dependence of *AI* of fibrinogen solution and fibrinogen solution supplemented either by thrombin or myeloperoxidase and thrombin. Arrows denote the moment of addition of the corresponding protein. (c) CD spectra of the solutions of fibrinogen and fibrinogen incubated with myeloperoxidase for 40 minutes. Fractions of protein secondary structure extracted from these data. Fg/MPO molar ratio is 21:1.

Top curve in Figure 4a represents an absorption spectrum of thrombin mediated fibrinogen clots obtained for the same fibrinogen concentration (150 µg/ml) in 30 minutes after thrombin addition. This spectrum is significantly higher than for fibrinogen–myeloperoxidase mixture indicating that thrombin-mediated fibrinogen clotting is more extensive than myeloperoxidase-induced fibrinogen clotting at the studied timescale (30 min). Moreover, the spectrum of fibrinogen–thrombin mixture is even higher than that of the fibrinogen–myeloperoxidase– thrombin mixture (third bottom curve). This fact may be explained by the decrease of the number of native fibrinogen molecules during fibrinogen incubation with myeloperoxidase prior to thrombin addition due to unfolding of a certain fraction of fibrinogen caused by its contact with myeloperoxidase (Figure 1d).

The *AI* value for fibrinogen solution, fibrinogen–myeloperoxidase mixture and fibrinogen– myeloperoxidase–thrombin mixture is plotted as a function of time in Figure 4b (red data points). Small *AI* values obtained for fibrinogen solution before myeloperoxidase addition (negative values on the timescale) indicate the absence of big protein aggregates. After myeloperoxidase addition *AI* significantly (by more than 10 units) increases during the next ∼19 minutes. This growth of *AI* may be rationalized by the formation of fibrinogen fibrils (Figure 2c and 3a) from the fraction of fibrinogen molecules unfolded by their contact with myeloperoxidase (Figure 1d). Further slight decrease of *AI* (1140 – 1940 s on the timescale) may be attributed to a precipitation of a small fraction of protein fibrils onto the walls of the optical cuvette that leads to a slight decrease of the number of protein fibrils in the bulk solution.

Addition of thrombin to the fibrinogen–myeloperoxidase mixture leads to a sharp jump of *AI* value by ∼20 units (1940 – 2050 s in the timescale) that is followed by its moderate increase by ∼10 units (2050 – 3700 s). The observed behavior of *AI* may be explained by the extensive formation of fibrils from the fraction of native fibrinogen present in fibrinogen–myeloperoxidase mixture via the conventional pathway of thrombin dependent fibrinogen clotting. For comparison, a time dependence of *AI* for thrombin mediated fibrinogen clotting (for the same fibrinogen concentration) is presented in Figure 4b (black line). *AI* of fibrinogen–thrombin system was higher than for fibrinogen–myeloperoxidase–thrombin system within 60 min of observation.

For estimation of protein secondary structure content in the fibrinogen–myeloperoxidase fibrils we have used CD. CD spectra recorded for a 40 min fibrinogen–myeloperoxidase mixture (at a fibrinogen concentration of 150 µg/ml) and for a pure fibrinogen solution of the same concentration were similar (Figure 4c). The contribution of myeloperoxidase to the CD spectrum is negligibly small due to comparably low myeloperoxidase concentration (Supplementary information, Figure S3). According to the obtained data, the content of secondary structured elements in fibrinogen solution is estimated as 27.1% and 24.3% for α-helix and β-strand, respectively (Figure 4c), being in general agreement with previously reported values ^36^. The content of secondary structured elements in a fibrinogen–myeloperoxidase mixture after 40 minutes of incubation was similar to that of fibrinogen (25.4% and 29.6% for α-helix and β-strand, respectively; Figure 4c) indicating that myeloperoxidase-induced fibrinogen clotting is not accompanied by significant change of the secondary structure. The similar conclusion has been made for a fibrinogen polymerization into fibrin after enzymatic cleavage by thrombin ^37^. Retaining of secondary structure in fibrinogen–myeloperoxidase clots resulted from CD spectra allows interpreting myeloperoxidase-induced fibrinogen unfolding (Figure 1d) as partial loss of fibrinogen tertiary rather than secondary structure.

### Effect of the ionic strength and pH on myeloperoxidase induced fibril formation

To study the effect of ionic strength (*I*) on myeloperoxidase induced fibril formation, we have recorded the light absorption spectra of a 1:1 (molar ratio) fibrinogen–myeloperoxidase mixture in three different media: water, 10 mM NaCl and 171 mM NaCl (pH 5.5). Immediately after addition of myeloperoxidase to fibrinogen solution a stepwise growth of *AI* (Δ*AI*_0min_) is observed in all three cases, however, it becomes less pronounced with the increase of the ionic strength (Figure 5a, closed symbols, the first and second data points in each dataset). Turbidity of fibrinogen solution in water right after addition of myeloperoxidase is so high that it can be confirmed visually (Figure 5c and Supplementary video). During the next 30 minutes the initial jump of *AI* is followed by considerable growth (Δ*AI*_30min_) for water or 10 mM NaCl solution but not for 171 mM NaCl solution, where *AI* remains, up to fluctuations, stable (Figure 2a, closed symbols, the third and following data points in each dataset). The Δ*AI*_0min_ (blue columns) and Δ*AI*_30min_ (red columns) values are summarized in the plot in Figure 5b (datasets 1 – 3). Grey columns in Figure 5b represent the *AI* growth obtained from the sum of the myeloperoxidase and fibrinogen spectra (Δ*AI*_sum_) recorded for the same conditions: this value represents a hypothetical *AI* growth of the fibrinogen–myeloperoxidase mixture on the assumption of the absence of their interaction. In water and 10 mM NaCl both Δ*AI*_0min_ and Δ*AI*_30min_ are significantly larger than Δ*AI*_sum_, indicating the myeloperoxidase induced fibrinogen aggregation. However, in 171 mM NaCl solution Δ*AI*_0min_ and Δ*AI*_30min_ are comparable with Δ*AI*_sum_. Therefore, the growth of *AI* in this case may be explained by an additive contribution of the two components of the mixture rather than myeloperoxidase induced fibrinogen aggregation, though the latter effect cannot be completely excluded.

**Figure 5.**
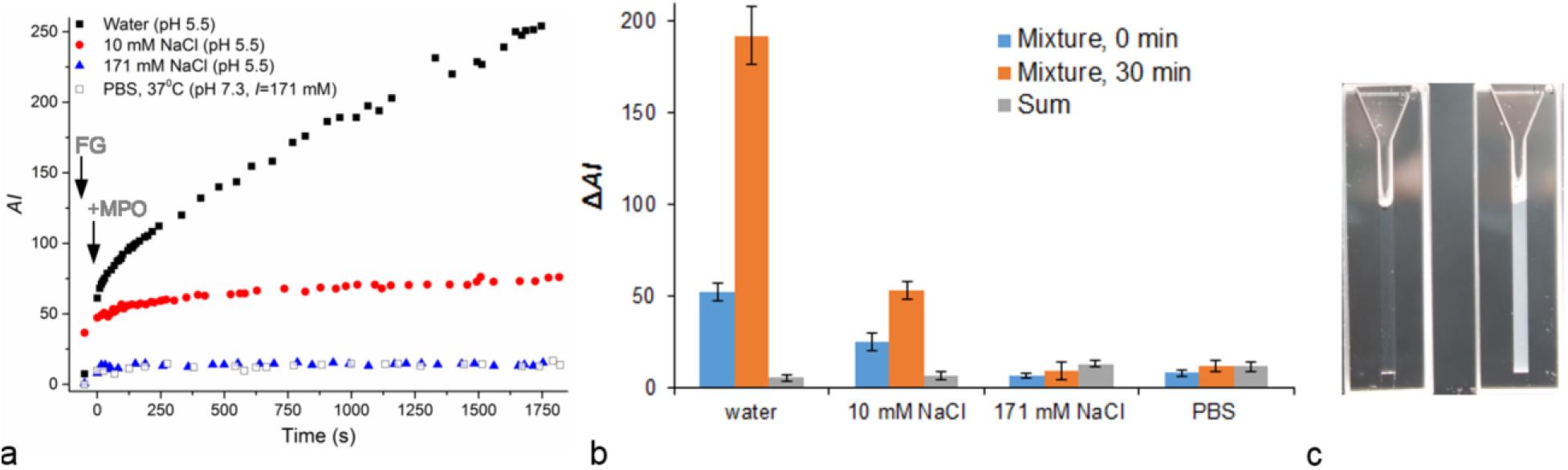
Fibrinogen fibril formation at different conditions. (a) *AI* values of fibrinogen solution before (first data point in each data set) and after (all other data points) addition of myeloperoxidase as a function of time (zero time corresponds to addition of myeloperoxidase) at pH 5.5 (water, 10 mM NaCl and 171 mM NaCl solutions; closed symbols) or at pH 7.3 (PBS solution; open symbols). (b) The change of *AI* (Δ*AI*) immediately and in 30 minutes after myeloperoxidase addition to fibrinogen solution as compared to Δ*AI* calculated for the sum of the myeloperoxidase and fibrinogen spectra obtained for the same conditions. Fg/MPO molar ratio is 1:1. (c) A photograph of the quartz cuvettes filled with water fibrinogen solution (left) or 2 minutes’ fibrinogen–myeloperoxidase mixture in water after short shaking (right).

The observed behavior of Δ*AI*_0min_ and Δ*AI*_30min_ at different ionic strengths can be interpreted by the Debye screening of electrostatic charges in electrolyte solutions. Since the Debye length is inversely proportional to the square root of the ionic strength of the solution, electrostatic attraction between fibrinogen and myeloperoxidase molecules is more impeded at higher ionic strengths. Therefore, myeloperoxidase induced fibrinogen clotting is also hampered upon growth of ionic strength of the solution, resulting in a smaller *AI* growth. Noteworthy, there is no detectable myeloperoxidase induced fibrinogen aggregation in 171 mM NaCl solution.

Finally, we have characterized light absorption spectra of a fibrinogen–myeloperoxidase mixture in PBS solution (pH 7.3, *I*=171 mM) at 37°C (Figure 5a, open symbols) that mimics physiological conditions. At pH 7.3 myeloperoxidase is still positively charged, whereas fibrinogen outer globules are even more negatively charged than at pH 5.5. Despite the opposite charges of the proteins, the obtained spectra do not reveal fibrinogen aggregation: the growth of *AI* is comparable to the value obtained for the sum of the of the myeloperoxidase and fibrinogen spectra (Figure 5b, the last dataset). This behavior of the *AI* is very similar to that observed at 171 mM NaCl solution (Figure 5a), which has the same ionic strength but more acidic pH (5.5).

Therefore, the ionic strength (and an associated Debye screening effect) has a greater effect on myeloperoxidase induced fibrinogen aggregation than pH associated changes of the proteins charges. In particular, the ionic strength of 171 mM (the Debye length ∼0.7 nm) prevents electrostatic attraction between fibrinogen and myeloperoxidase molecules independent of the values of the opposite charges of the proteins (within the studied pH range), whereas at the ionic strength ≤10 mM (the Debye length ≥3 nm) the protein electrostatic attraction becomes possible. Strong dependence of the myeloperoxidase induced fibrinogen clotting on the ionic strength represents a convenient tool for regulation of this process, which may be used for the development of fibrinogen based matrixes.

The ability of fibrinogen to form fibers in vitro in the absence of thrombin has been widely reported and associated with the effects of a substrate surface and drying, which can be accompanied by one of the following factors: denaturing buffers (such as organic solvents) ^38^, highly acidic pH (<4) ^39^, elevated temperatures ^26^, high fibrinogen concentrations (>2 mg/ml), high ionic strength (up to 0.75M) ^36,40^, shear forces ^41^. In contrast, our results evidence a novel way of inducing fibrinogen fibril formation, since it proceeds in a bulk solution at relatively low fibrinogen concentrations (<0.1 mg/ml), zero (or low) ionic strength and in absence of any organic solvents (even in water). Due to non-specific, electrostatic origin of the revealed fibrinogen fibril formation we assume that myeloperoxidase is not a unique protein inducing fibrinogen unfolding and extensive fibril formation at low ionic strength. There is a big potential of utilization of other proteins to induce and regulate thrombin free fibrinogen fibril formation for the development of new fibrinogen based materials with tuned properties.

### Conclusions

By the example of myeloperoxidase-induced fibrinogen unfolding we have revealed that ATP independent, non-targeted protein unfolding may be mediated by contact with another protein. Due to high biocompatibility and nontoxicity of the proteins, their use as a factor that induces protein unfolding may have a particular interest in biotechnology and medicine.

Electrostatics was shown to play a primary role in fibrinogen complexation with myeloperoxidase and further fibrinogen unfolding. Fibrinogen molecules unfolded by myeloperoxidase tend to form fibrils and clots that differ from those formed by fibrin after fibrinogen cleavage by thrombin. Non-conventional (i.e. not mediated by thrombin) fibrinogen clotting decreases the number of intact fibrinogen molecules, which retain the ability to conventional (i.e. thrombin mediated) clotting. Myeloperoxidase-induced fibrinogen clotting is not detected in the physiological buffer though may not be totally excluded. This effect becomes pronounced at low ionic strengths (0–10 mM) indicating the important role of Debye screening in regulation of fibrinogen fibril formation. Due to non-specific character of fibrinogen– myeloperoxidase interaction, other proteins with similar effects may be found in future. The revealed effects may be potentially used in biomedical applications, such as development of fibrinogen based matrices with tuned properties and regulation of fibrinogen clotting and platelet adhesion.

## Methods

### AFM

GM-HOPG surface was prepared by application of 10 μl of 10 µg/ml GM (Nanotuning, Russia) solution in double-distilled water on a freshly cleaved HOPG (ZYB quality, NT-MDT, Russia) surface for 10 s followed by the removal of the droplet from the surface by a nitrogen flow. Fibrinogen (Calbiochem, Germany) water solution was mixed with either myeloperoxidase (Planta Natural Products, Austria) or horse spleen ferritin (Sigma-Aldrich) water solution so that final concentration of each component in the mixture was 1 μg/ml and the total volume 2 µl. The resulting fibrinogen–myeloperoxidase or fibrinogen–ferritin mixture was incubated for 3 s – 10 min prior to deposition on a surface. In some experiments fibrinogen solution (1 μg/ml) or 1 min fibrinogen–myeloperoxidase mixture was supplemented by thrombin solution so that its final concentration was 0.2 μg/ml.

For AFM investigations 0.5 μl of a protein mixture (fibrinogen–myeloperoxidase, fibrinogen– ferritin, fibrinogen–thrombin or fibrinogen–myeloperoxidase–thrombin) prepared as described above was deposited onto a GM-HOPG surface for 5 s. After that, 100 μl of water was added on the surface (rinsing) followed by immediate removal of the whole droplet by a nitrogen flow.

AFM experiments were performed on the multimode atomic force microscope Ntegra Prima (NT-MDT, Russia) operated in an ambient environment in an attraction regime of an intermittent contact mode with a cantilever oscillation amplitude of ∼1 nm. Ultra-sharp tips (carbon nanowhiskers with a curvature radius of several nanometers grown at tips of commercially available silicon cantilevers with a spring constant of 5–30 N/m) were used. The scan rate was typically 1 Hz with 1024×1024 pixels. The pixel resolution was 1 pixel/nm that provides high accuracy of the determination of the mean height of protein globules (pixelization associated error was less than 2% for the studied proteins)^42^.

### Absorbance spectrophotometry

In the experiments performed in water 1 cm quartz cuvette (Bio-Rad Labs, USA) was filled with 300 µl of fibrinogen water solution (150 µg/ml) and placed to the NanoDrop Spectrophotometer (Thermo Scientific, USA). After recording of several absorption spectra of fibrinogen solution, 0.5 µl of myeloperoxidase or thrombin water solution was added to the cuvette (so that final concentration of myeloperoxidase or thrombin was 3 µg/ml that corresponds to ∼21:1 fibrinogen: myeloperoxidase molar ratio), and successive absorption spectra of a fibrinogen–myeloperoxidase or fibrinogen–thrombin mixture were recorded. In 30 minutes after myeloperoxidase addition the fibrinogen–myeloperoxidase mixture was supplemented with 0.5 µl of thrombin water solution (so that final concentration of thrombin was 3 µg/ml).

For investigation of the effect of ionic strength and pH, the quartz cuvette was filled with 300 µl of water or NaCl/PBS solution of fibrinogen (71 µg/ml) and placed to the spectrophotometer. After recording of several absorption spectra of fibrinogen solution, 5 µl of water or NaCl solution of myeloperoxidase were added to the cuvette (so that final concentration of myeloperoxidase was 28 µg/ml that corresponds to ∼1:1 fibrinogen–myeloperoxidase molar ratio) and successive absorption spectra of a fibrinogen–myeloperoxidase mixture were recorded. The sum of the adsorption spectra of 71 µg/ml fibrinogen solution in water or NaCl/PBS and 28 µg/ml myeloperoxidase solution in the same medium (recorded separately) was used as the controls.

For quantitative characterization of fibrinogen fibril formation we have used an aggregation index (*AI*) ^43^:

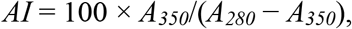

where *A*_*280*_ and *A*_*350*_ are the measured absorbance at 280 and 350 nm, respectively.

The spectra in water and NaCl solutions were recorded at 21°C; in PBS solution – at 37°C. The standard deviations obtained from three independent experiments were used as errors for *AI* values.

### SEM

Preparation of fibrinogen–myeloperoxidase, fibrinogen–thrombin or fibrinogen– myeloperoxidase–thrombin mixtures for SEM analysis was the same as for spectrophotometric experiments in water. The time of incubation of fibrinogen with myeloperoxidase or thrombin or a fibrinogen–myeloperoxidase mixture with thrombin was 1 min. A ∼2 mm stripe of HOPG with a freshly cleaved surface was immersed into a protein mixture for 10 s and left in ambient atmosphere until complete drying.

SEM experiments were performed on the Zeiss Merlin microscope equipped with GEMINI II Electron Optics (Zeiss, Germany) with accelerating voltage 1 kV and probe current 90–100 pA using high efficiency secondary electron (HE-SE2) detector.

### Circular dichroism spectroscopy

200 µl of either 150 µg/ml fibrinogen water solution or fibrinogen (150 µg/ml)– myeloperoxidase (3 µg/ml) mixture was loaded in the narrow quartz cuvette with an optical path of 0.5 mm. CD spectra were obtained in a Chirascan spectrophotometer (Applied Photophysics, UK) at 21°C. Spectra deconvolution was performed using a BeStSel web server (http://bestsel.elte.hu) ^44,45^. The relative error of the protein secondary fraction determination was 2%.

## Acknowledgments

We thank Dr. A.V. Sokolov (Institute of Experimental Medicine, Saint Petersburg, Russia) for preparation of myeloperoxidase and the valuable consultations, and Dr. I. I. Vlasova (I.M. Sechenov Moscow State Medical University, Moscow, Russia) – for the valuable consultations. This work was supported by the Russian Science Foundation [17-75-30064 to D.V.K.]. E.V.D. acknowledges the financial support by the Ministry of Education and Science of the Russian Federation in the framework of increase Competitiveness Program of NUST “MISIS”, implemented by a governmental decree dated 16th of March 2013, No 211 (electrostatic surface potential estimation).

