## Supplementary information for "Myeloperoxidase-induced fibrinogen unfolding and clotting"

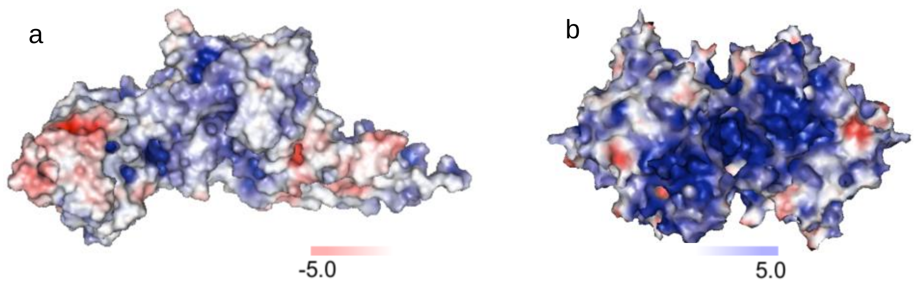


**Figure S1.** Molecular surfaces of (a) fibrinogen outer globular domain (PDB entry 3HUS) and (b) myeloperoxidase (PDB entry 1MHL). Colors refer to localized electrostatic potential at pH 5.5: blue – positive, red – negative, white – neutral. Surface potentials were built using APBS/PDB2PQR software [1,2].

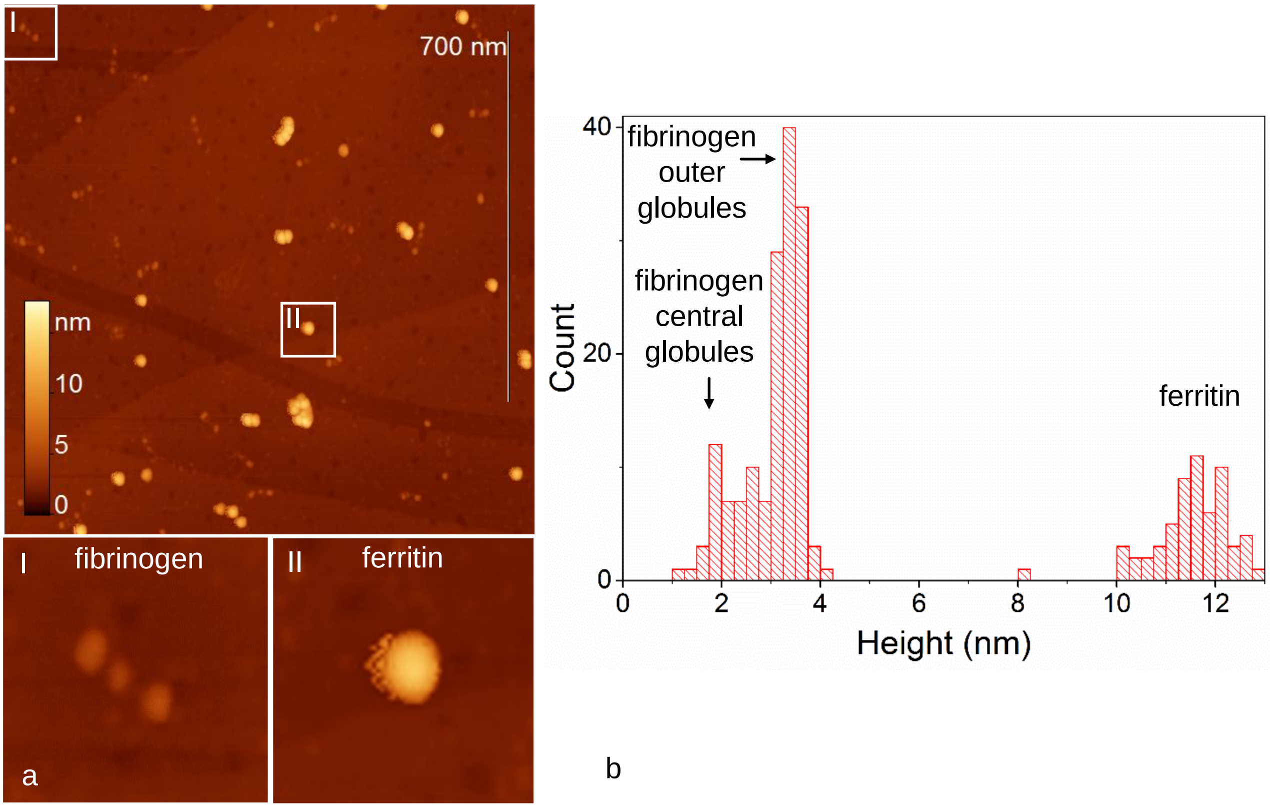


**Figure S2.** (a) AFM image of a fibrinogen–ferritin mixture incubatred for 30 min, deposited onto a GM-HOPG surface. The regions marked with a box and labelled with Roman numbers are enlarged at the bottom. The sizes of the images are 1000 × 1000 nm$^2$ (enlarged regions 100 × 100 nm$^2$). (b) Height distribution histogram of the objects observed in the AFM images of a fibrinogen–ferritin mixture.

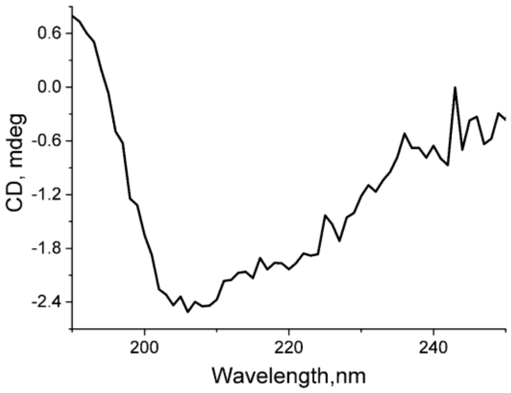


**Figure S3.** A typical CD spectrum of 6 µg/ml MPO solution (this concentration is 2 times higher than final MPO concentration in the FG–MPO mixtures).
